# Looking for the needle in a downsized haystack: Whole-exome sequencing unravels how selection and gene flow have shaped climatic adaptation in Douglas-fir (*Pseudotsuga menziesii*)

**DOI:** 10.1101/2020.11.16.381004

**Authors:** Jan-Peter George, Silvio Schueler, Michael Grabner, Sandra Karanitsch-Ackerl, Konrad Mayer, Michael Stierschneider, Lambert Weissenbacher, Marcela van Loo

## Abstract

The widespread Douglas-fir (*Pseudotsuga menziesii (*Mirb.) Franco) occurs along a steep gradient of diverse climates throughout its natural range, which is expected to result in spatially varying selection to local climate conditions. However, phenotypic signals of climatic adaptation can often be confounded, because unraveled clines covary with signals caused by neutral evolutionary processes such as gene flow and genetic drift. Here, we present phenotypic and genotypic data from a common garden experiment showing a putative signal of adaptation to climate after trees have been growing for 40 years in a common environment. Sixteen Douglas-fir provenances originating from a North-to-South gradient of approx. 1,000 km were analyzed and genomic information was obtained from exome capture, which resulted in an initial genomic dataset of >90,000 single nucleotide polymorphisms. We used a restrictive and conservative filtering approach which permitted us to include only SNPs and individuals in environmental association analysis (EAA) that were free of potentially confounding effects (LD, relatedness among trees, heterozygosity deficiency and deviations from Hardy-Weinberg proportions). We used four conceptually different genome scan methods based on F_ST_ outlier detection and gene-environment association in order to disentangle truly adaptive SNPs from neutral SNPs and found that a relatively small proportion of the exome showed a truely adaptive signal (0.01-0.17%) when population substructuring and multiple testing was accounted for. Nevertheless, the unraveled SNP candidates showed significant relationship with climate at provenance origins which strongly suggests that they have most likely featured adaption in Douglas-fir across a steep climatic gradient. Two SNPs were independently found by three of the employed algorithms and one could be assigned with high probability to a *Picea abies* homolog gene involved in circadian clock control as was also found in *Populus balsamifera*.

## 1. Introduction

Conifers have succesfully occupied a large number of habitats and different climates throughout their past-glacial histories, which allowed them to survive and establish in harsh environments but also to colonize ecoregions with high productivity and optimal growing conditions (Farjon, 2010). As such, Douglas-fir (*Pseudotsuga menziesii*) constitutes one of the most widespread conifers in western North America with a distribution from Southern California up to Northern British Columbia (Gugger et al. 2010). Within its current range of occurrence two distinct varities are known which appear genetically and phenotypically different: the coastal variety (also called *P. menziesii* var. *menziesii*) and the interior variety (*P. menziesii* var.*glauca)*. While the coastal variety is mainly found from central California to coastal British Columbia along the Pacific coast, the interior variety has its main south-to-north expansion from Wyoming/Montana over Alberta up to central British Columbia (Hermann & Lavender 1990; Martinez, 1949). Both of these macro-geographic regions differ substantially in climate with cool and moist climate conditions in the Pacific habitats towards more continental and dry growing conditions in the interior (Gugger et al. 2010). Both habitats are separated by the Great Basin and Columbia Plateau, respectively. Accordingly, several common garden experiments found intra-specific trait differences among Douglas-fir provenances that were related to growth, physiology, and phenology and which showed associations with differences in source climate among provenance locations (e.g. Anekonda et al. 2004; Fielder & Owens 1989; Kleiber et al. 2017; Pharis & Ferrel 1966). While such trait-climate associations can indeed suggest patterns of local adaptation due to spatially varying selection (Leinonen et al. 2013), their interpretation can still be doubtful, because the climatic clines at which provenances occur often show parallelism with re-colonization routes after population contraction and expansion (Nadeau et al. 2016). Consequently, the putative adaptive signal obtained from such trait analyses can be confounded by those that were left behind by neutral processes such as genetic drift and gene flow, which have often resulted in strong population substructuring (Günther & Coop, 2013).

Incorporating information from genetic markers has significantly improved our understanding of the role of neutral processes in population structuring in many plant species and has shed light into their postglacial migration histories (Hewitt, 1999). However, the main challenge still remained, which is to distuingish truly adaptive regions in the genome from those which show strong divergence just as a result of neutral genetic processes. Identifying true environmental associations and adaptive regions in the genome (e.g. from <u>s</u>ingle <u>n</u>ucleotide <u>p</u>olymorphisms or SNPs) is of utmost importance, since only adaptive markers have the potential to assist tree breeding of more resistent genotypes and selection of climatically adapted genotypes for forest management under climate change (e.g. Grattapaglia et al. 2018; Harfouche et al. 2012; Neale & Kremer, 2011). Yet, one of the strongest limitations in disentangling adaptive from neutral genomic locations is to aquire a marker set which is large and dense enough to cover the genome in a representative and unbiased fashion. This is particularly difficult in conifers such as Douglas-fir, which harbour huge genomes in the magnitude of several gigabases (Neale et al. 2017; Nystedt et al. 2013; De La Torre et al. 2014). Nevertheless, recent improvement in massive parallele sequencing and bioinformatics have made it possible to reduce the complexity of such genomes by limiting it to the protein coding region, which is called the *exome* (e.g. Neves et al. 2013). Exome capture in conifers results in massive complexity reduction which allows to rapidly generate markers at high number and relatively low costs in order to investigate genomic regions of interest thoroughly (see for instance Capblancq et al. 2019; Suren et al. 2016; Thistlethwaite et al. 2017 for examples on conifers). Even though the exome represents only a small fraction of the total genome, this data is particularly suited for environmental-association-analyses (EAA), since adaptive candidate SNPs can be further interrogated when a gene model or annotated reference genome is avialable. Moreover, by using a subset of SNPs in the exome which show no significant deviation from neutrality expectation (e.g. F_st_), the dataset can be used for estimating population substructuring as well.

In this study, we investigate a 40-year old common garden experiment with Douglas-fir provenances from 16 accessions across its distribution by combining dendroclimatological methods with modern sequencing technique which allowed us to generate a dense set of single nucleotide polymorphisms throughout the entire exome. Based on the inital finding that provenances showed a strong association between growth traits (response of annual increment to summer temperature) and seed source climate, we used classical F_ST_-outlier approaches, bayesian inference methods, and latent factor mixed modelling (LFMM) in order to identify SNPs with a,, truely” adaptive signal for climate adaptation. We hypothesize that spatially varying selection has shaped actual patterns of trait differentiation in Douglas-fir despite a high amount of neutral covariance caused by genetic drift and gene flow.

## 2. Materials &Methods

### 2.1 Common garden experiment

The analyzed trees originate from a block-replicated common garden experiment that was established in 1976 in eastern Austria with a total of 49 native Douglas-fir seed sources. The mean annual temperature of the trial site is 7.4°C and mean annual precipitation is 650mm (average over the period from 1961-1990; source: Austrian Central Station for Meteorology). Trees have been planted as two-year-old plants in a 2×2m spacing and each provenance was replicated three times in a random fashion within the trial. In 2012, two cores per tree were taken from a subset of 16 provenances originating from a wide part of the natural distribution encompassing Oregon, Washington and British Columbia (see Fig. 1, Tab. 1 and more information below). In brief, tree cores were taken at breast height for a total of 178 trees (9-15 per provenance and evenly sampled across the three blocks) and stored in plastic tubes until further processing in the lab. Cores were then cut into approx. 1.4mm wide cross sections with a double-blade circular saw, placed on microfilms and exposed to a 10 kV (24mA) X-ray source for 25 minutes. Thereafter, the obtained microfilms were analyzed using the software WinDENDRO 2009 (Regent Instrument, Quebec, Canada) by measuring annual increments to the nearest 0.001 mm for early- and latewood, respectively (see also George et al. 2019, 2017, 2015 for more information on the methodolgy). The 50% percentile of the density distribution between minimum and maximum density of each ring was used for the identification of earlywood-latewood boundaries (Fries & Ericsson 2009; illustrated in Supporting Information Fig. S1). The radial increment data from this common garden experiment, which is in detail described in George et al. (2019) was re-analyzed for this study and therefore additional needle samples for DNA analyses were taken from the same 178 trees in May 2019. Needles were shot from the lower part of the tree crowns by using a shot gun and stored in silica gel until DNA extraction.

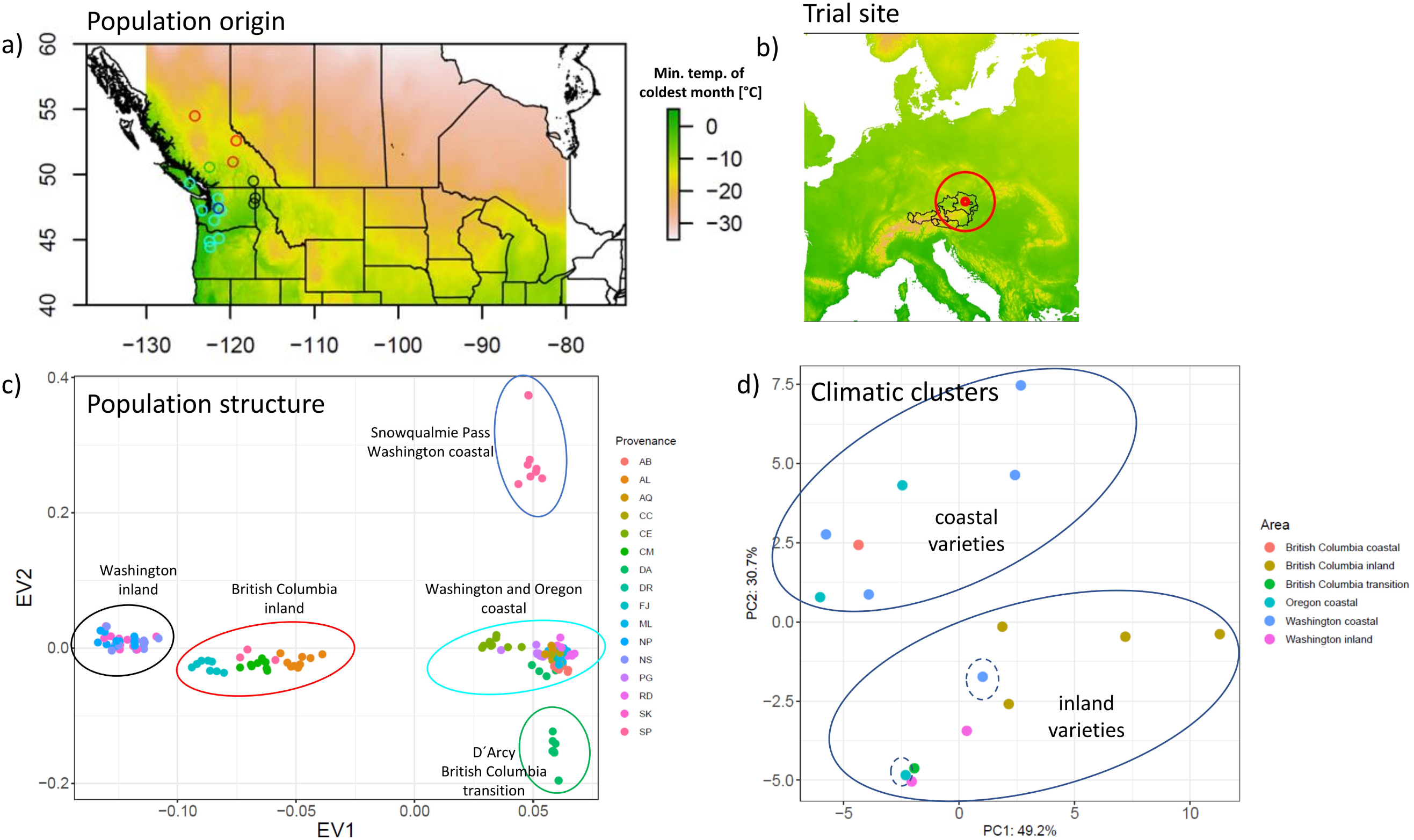

### 2.2 DNA analysis, probe design & SNP calling

For each sample, genomic DNA was extracted from lyophilized needle tissue using a CTAB protocol (van der Beek et al. 1992) with minor modifications made for the processing of 96-well plates. DNA concentration and quality was assessed with Qubit® QuantIT dsDNA BR Assay kit and a Qubit1 Fluorometer (Thermofisher, Waltham, MA, USA). A total volume of 50 µL genomic DNA for each sample (average concentration: 24 ng/ µL) was sent to the genotyping service provider RAPiD Genomics LLC (Gainesville, FL, USA) for sequencing. Exome capture probes were designed as described in Neves et al. (2013): briefly, DNA was sheared to a mean fragment length of 400bp, fragments were end-repaired, followed by incorporation of unique dual-indexed Illumina adapters and PCR enrichment. Intron-exon boundaries were designed by mapping transcripts from the published Douglas-fir reference transcriptome from Howe et al. (2013) (https://www.ncbi.nlm.nih.gov/Traces/wgs/?val=GAEK01) to the Douglas-fir reference genome assembly *Dougfir 1*.*0* (https://www.ncbi.nlm.nih.gov/assembly/GCA_001517045.1/#/st) from Neale et al. (2017). Only probes that hit once to the genome and which showed sufficient GC content were included. This resulted in 20K probes that were found in 12,272 scaffolds with an average number of 1.6 probes per scaffold and a maximum number of 18 probes per scaffold. Sequence capture was performed using RAPiD Genomics proprietary chemistry and workflows. Samples were pooled equimolar and sequenced using a HiSeq 2×150.

After sequencing, *Trimmomatic* (Bolger et al. 2014) was used to remove sequencing adapters and the trimmed reads were mapped with BWA (Li et al. 2009) using the default settings to the Douglas-fir reference genome. SNPs within probes were identified with *FreeBayes* (Garrison & Marth, 2012, available at https://github.com/ekg/freebayes). *VCFtools* (Danacek et al. 2011) was used to store the identified SNPs for further downstream analysis in VCF format by applying the following filter: accepted mean depth across all individuals=750, minimum total depth per individual=3, Percent of individuals allowed missing to retain a site=0.4, minimum SNP quality=10.

### 2.3 Consecutive SNP filtering and exclusion of trees with confounding effects

In order to create a genomic dataset that is free of sources which could potentially affect the false positive discovery rate of adaptive SNPs, we used a restrictive and conservative filtering approach: as such, we removed all SNPs with a minor allele frequency<0.05 as well as those SNPs that showed significant deviation from Hardy-Weinberg expectation (threshold: 10^−6^). Furthermore, we performed pairwise LD pruning between markers with a LD threshold of 0.2 in order to retain only unlinked SNPs for environmental association analysis. For this, we used the *snpgdsLDpruning* function implemented in the *SNPrelate* package in R (R Development Core Team 2003).

Furthermore, kinship among individuals was estimated by calculating the identity-by-descent (IBD) methods of moments between all pairs of individuals using a subset of SNPs that was already corrected for linkage disequilibrium (see section above). We retained only those trees that showed a kinship coefficient <0.25 to any other tree in the dataset. Trees that showed signals of heterozygote deficit (calculated as 1-(Het_obs_/Het_Exp_) over all loci) were removed prior to analysis with a threshold of 0.1. Finally, we included only trees with an overall SNP call rate of 0.95 for subsequent analyses. All filtering steps were performed in R by using the packages *gdsfmt, SNPrelate* (Zheng et al. 2012), and *vcfR* (Knaus & Grünwald, 2017).

### 2.4 Climate data & trait analyses

Information on provenance climate was obtained from the ClimateNA database (Wang et al 2016) under http://www.climatewna.com/. Latitude/longitude and elevation information for provenance origins were used to retrieve a total of 247 annual, seasonal, and monthly climate variables (reference period: 1961-1990). The full list of climate variables can be found in the **Supplementary Information S1**. In order to reduce the complexity of this data, the 247 variables were transformed into principal components and the first four PCs were standardized (i.e. expressed as standard deviations from the mean). These standardized climate PCs (hereafter called environmental PCs) were used for subsequent analyses of environmental association analysis.

Growth traits for this dataset were derived from the study in George et al. (2019). Briefly, tree-ring series were standardized using a 15-year cubic smoothing spline in order to remove the biological age trend. The standardized series were aggregated to chronologies for each provenance by calculating the year-to-year biweight robust mean among trees in each provenance (Bunn 2008). For the purposes of this study we used the bootstrapped response functions that were applied to ring width chronologies of the same Douglas-fir provenances in George et al. (2019). Response functions and in particular response coefficients between a growth trait (in our case ring width) and trial site climate provide insights of the relative importance of an environmental variable for a trait (e.g. Levesque et al. 2014; Zang & Biondi 2013). However, when calculated at provenance level, bootstrapped response coefficients can be related to provenance source climate in order to unravel genecological clines (that is: some provenances may react more sensitive to an environmental factor than others as a result of local adaptation). For environmental association analysis in this study we used the response of earlywood increment to July temperature at trial site because of two reasons: first, the data in George et al. (2019) showed significant differentiation among Douglas-fir provenances for this trait and, second, growth response in earlywood has been shown to be significantly involved in drought response and subsequent survival in Douglas-fir (e.g. Martinez-Meier et al. 2008) which suggests that this trait most likely harbours signals of spatially varying selection among provenances. Multi-year correlations between standardized earlywood time series and July temperature at trial site between 1979 and 2011 were bootstrapped by using the R packages *bootRes* (Zang & Biondi, 2013) and *dplR* (Bunn 2008).

### 2.5 Outlier detection & environmental association analysis (EAA)

The aim of this study is to distuingish SNP markers throughout the exome which have a putative adaptive signal for an environmental factor from non-adaptive SNPs. For this, we used four different algorithms which are either based on F_ST_ outlier detection methods (Arlequin & BayScEnv) or on correlations between allele frequencies and environmental variables (BayEnv2 & LFMM2). Regardless of the algorithm that is used for doing this, controlling for variation in allele frequencies that is simply caused by neutral evolutionary processes such as genetic drift or gene flow is a crucial step for identifying adaptive candidate SNPs (Rellstab et al. 2015). Since the fraction of such adaptive SNPs is usually small compared to all analyzed SNPs, adjusting the results carefully for false positive discovery rates is of utmost importance. We therefore picked four up-to-date population genomic programs (Arlequin, BayScEnv, BayEnv2, LFMM2) which have implemented various testing strategies in order to control for confounding effects and false positive errors.

The coalescent approach implemented in *Arlequin* 3.5 (Excoffier & Lischer 2010) compares locus-specific patterns of population differentiation to a null distribution which is simulated under the assumption of a hierachical island model (Excoffier et al. 2009). P-values are estimated locus-wise from the joint distribution of heterozygosity and F_ST_. If the locus-specific F_ST_ exceeeds the confidence boundaries of the global observed F_ST_ the respective locus is considered to be an outlier and likely to be under selection. Since we observed significant population substructure (see results below) in our dataset, we chose the hierachical island model instead of the finite island model in order to reduce the number of false positive candidate SNPs, since confidence intervals around a given F_ST_ value become narrower the more populations are sampled (see Excoffier et al. 2009). Groups for the hierachical island models were defined based on the first two principal components obtained from 1,500 randomly chosen SNPs. We used 200,000 simulations in Arlequin vers. 3.5 with 10 simulated groups and 100 demes in each group. SNPs where considered to be under selection when their observed F_ST_ exceeded the 5% upper quantile of the simulated null distribution.

BayScEnv (Villemereuil & Gaggiotti, 2015) is a F_st_-based genome-scan method that incorporates also environmental differentiation. The algorithm tests a model of local adaptation with environmental differentiation (i.e. the parameter **g** in Villemereuil & Gaggiotti, 2015) against two alternative models: i) the neutral model consisting largely of demographic effects and ii) the locus-specific model which includes locus-specific effects other than selection due to environmental differentiation. Posterior error probabilities for chosing the model of local adaptation in favour of the two alternative models are calculated by means of MCMC simulations and results are also adjusted for false discovery rates by providing q-values for each locus. We used 20 pilot runs each with a length of 2000 and a burn-in of 50,000 with a thinning interval of 10. Furthermore, a prior probability of 0.1 was used for the parameter *pi* (assuming every 10th locus harbours a non-neutral signal), while a prior probability of 0.5 was used for the parameter p (assuming around 50% of all loci are associated with an environmental variable). Acceptance rates and convergence of the MCMC chain were inspected after each model run and for each of the four environmental variables (i.e. the standardized climate PCs) by using the *coda* package in R.

In contrast to the F_ST_-based outlier detection method *Bayenv2*.*0* (Günther & Coop 2013) calculates standardized allele frequencies and tests for correlations between environmental variables and those frequencies. Neutral and spurious correlations arising from shared population history and gene flow are controlled for by including a covariance matrix obtained from a subset of unlinked and putatively neutral SNPs. We used a subset of 1,500 randomly chosen unlinked SNPs for matrix estimation and ran Bayenv2.0 with 100,000 iterations. Subsequently, all identified SNPs for each of the four environmental variables with a Bayes factor > 30 (Jeffreys 1961) were considered as candidates for environmental association analysis. Since the algorithm in Bayenv2.0 is based on MCMC simulations, each run was repeated several times in order to ensure that results remain robust. We additionally calculated spearman rank correlation coefficients in addition to Bayes factors for ach SNP in order to ensure that detected candidates were correctly identified and not confounded by outliers (see recommendation in Günther & Coop, 2013).

Finally, we used a regression model combining fixed (i.e. environmental) and latent model effects (i.e. neutral population structure). The model has the form:

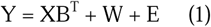

With Y being the response genotype matrix, XB^T^ the matrix of fixed effects (climate PCs), E the matrix of residual errors, and W the „latent matrix” (see Caye et al. 2019). W determines the confounding effect of neutral population structure and is determined by k latent factors, where k defines the number of ancestral populations as obtained from a subset of putative neutral loci. As an approximation of k, we used the screeplot obtained from principal component analysis performed on the 1,500 randomly chosen SNPs described above. Calculations were carried out using the *lfmm2* function in the LEA package in R (Frichot & Francois, 2015).

### 2.6 Adaptive signal of outlier SNPs and functional interrogation

In order to further interrogate the identified candidate SNPs, we first defined a set of consensus variants. Consensus SNPs were defined as those SNPs which were independently detected by at least two of the four chosen methods according to thresholds in Tab. 2. From those SNPs 500 bp long fragments were extracted (250 downstream and 250 bp upstream of the variant) and blasted against the Norway spruce (*Picea abies*) genome assembly (*Picea abies* 1.0 (complete) available under www.congenie.org) since this is so far the closest relative to Douglas-fir having an functionally annotated genome assembly. For every homolog contig that was found E-values, average identity, and putative gene function (if known) were reported. We always selected the homolog with the lowest E-value (i.e. number of hits one can expect by chance alone) and -if more than one gene was known to be located on that homolog-all putative gene functions were reported.

Finally and in order to prove whether the unraveled consensus SNPs have the ability to discriminate between provenances adapted to different climates, the first and second eigenvectors for each tree were calculated based on consensus SNPs and, for comparison, based on an equal number of randomly chosen neutral SNPs, respectively. For this, the *snpgdsPCA* function from the SNPRelate package in R was employed.

## 3. Results

### 3.1 SNP dataset

Within the 20K probes which were designed for exome capture, a total of 90,979 SNPs were successfully called with *Freebayes* and passed the applied quality filter as described in section 2.2. Since a large proportion of these 90,979 SNPs had either a minor allel frequency <0.05 or a call rate <0.95, 41,009 SNPs were consequently removed and not considered for downstream analysis. 4,269 markers showed significant deviations from Hardy-Weinberg proportion and were excluded from analyses accordingly. Finally, testing for linkage disequilibrium among markers revealed 17,489 SNPs which were physically unlinked and only these SNPs were included in all subsequent analysis steps.

From the 178 trees, 5 showed significant heterozygosity deficiency probably as a result of inbreeding and were hence excluded from subsequent analyses. There was no significant family structure among trees within provenances as revealed by IBD methods of moments so that all remaining 173 trees were included in environmental association analysis.

### 3.2 Climatic groups, population structure & growth response functions

The first two principal components of the 250 long-term climatic variables together explained 79.9% of variation (Fig. 1). The first principal component (hereafter called climate PC1) was strongly related to temperature variables (correlation to mean annual temp. −0.99 and to minimum temperature of coldest month: −0.97) and length of growing season (correlation to number of growing degree days>5°C=0.88). The second component (climate PC2) was strongly correlated with precipitation regime at seed origin (r_MAP_=0.94; r_summer:heat mositure index_=0.81). The third and fourth PCs explained less variation and were related to solar radiation and annual snow precipitation, respectively. As expected, provenances could be roughly dividend into two main groups when plotted against the first two climate PCs, that is inland and coastal provenances (Fig. 1d). Provenances Cle Elum (WA) and Pine Grove (OR) clustered climatically more closely with inland varieties, although they belong geographically still to the coastal zone.

Based on the subset of 1,500 randomly chosen SNPs five population clusters were unraveled: i) provenances from northern inland British Columbia (Fort St. James, Clemina, Adams Lake), ii) the three inland varieties from interior Washington and Southern British Columbia (Newport, Spokane (both WA) and Nelson (BC)), iii) coastal varieties from British Columbia, Washington and Oregon (Alberni, Matlock, Cle Elum, Pine Grove, Randle, Darrington, Abiqua Basin & Cascadia), iv) a single provenance cluster Snowqualmie Pass (WA) and v) another single provenance cluster including provenance D’Arcy in the transition zone between coastal and inland British Columbia (Fig. 1c).

Response of earlywood growth to July temperature at the trial site varied significantly among provenances. In general, inland provenances from British Columbia and Washington had lower response coefficients than their coastal counterparts (Fig. 2). Response coefficients varied between 0.17 (Newport WA) to 0.42 (Cascadia OR). Differences among provenances were statistically significant as suggested by 95% confidence intervals of the bootstrapped coefficients. When regressed against climate PC1 and PC2 the relationship was significant with a stronger positive earlywood response towards warmer provenance origins (r=0.57; Fig 2a) as well as towards wetter provenance orgins (r=0.48; Fig 2b). There were no significant relationships to climate PC3 and climate PC4 and consequently only the first two climate PCs were considered for environmental association analysis.

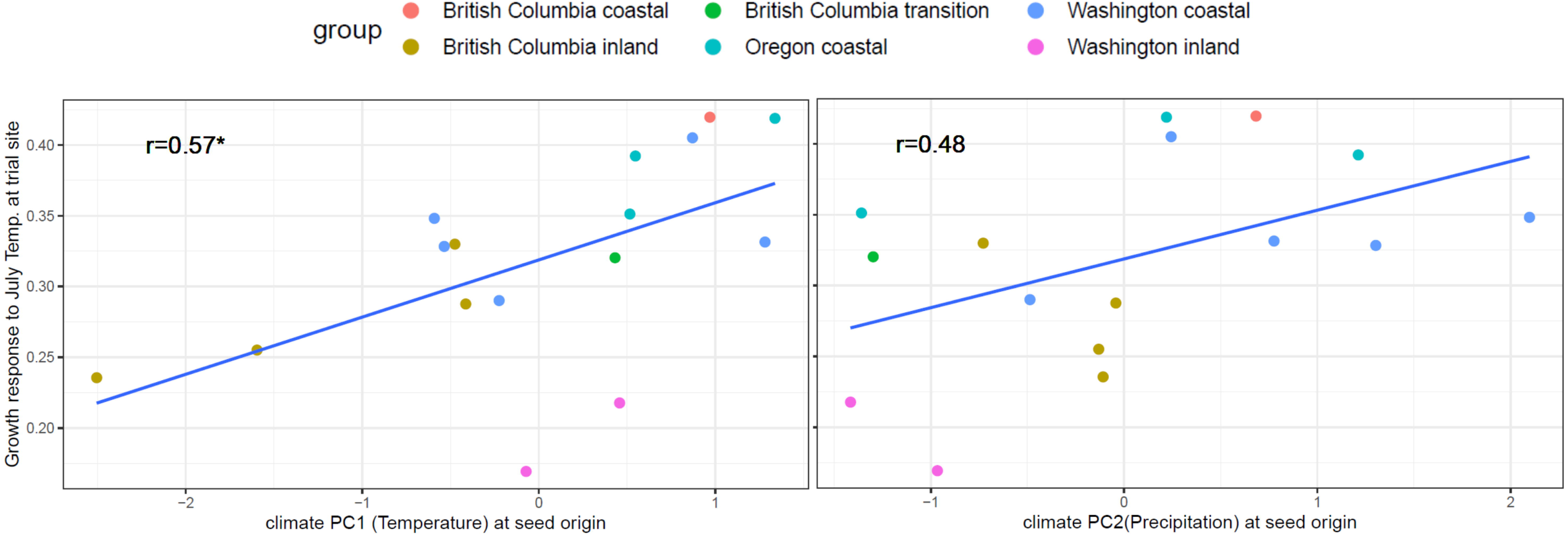

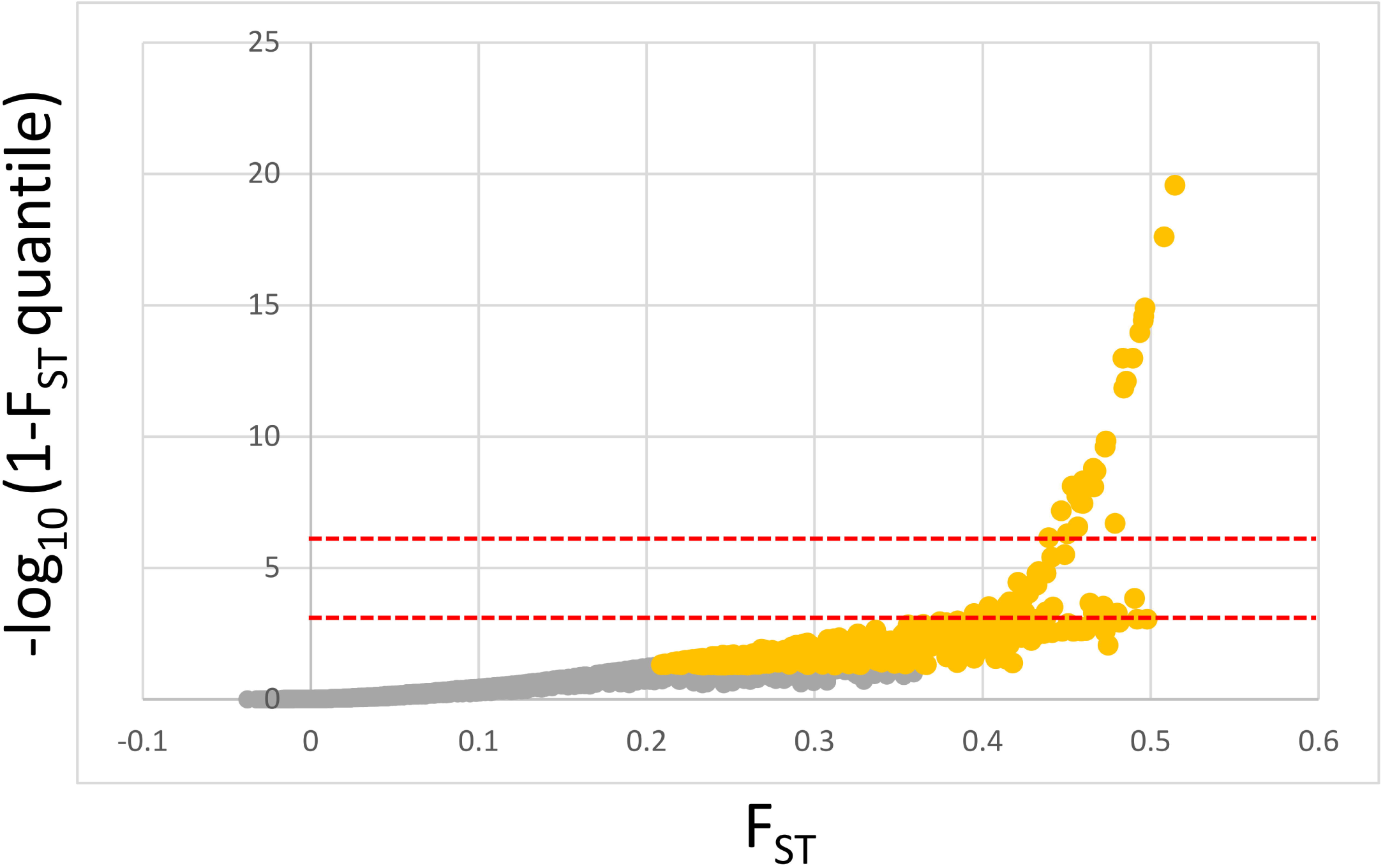

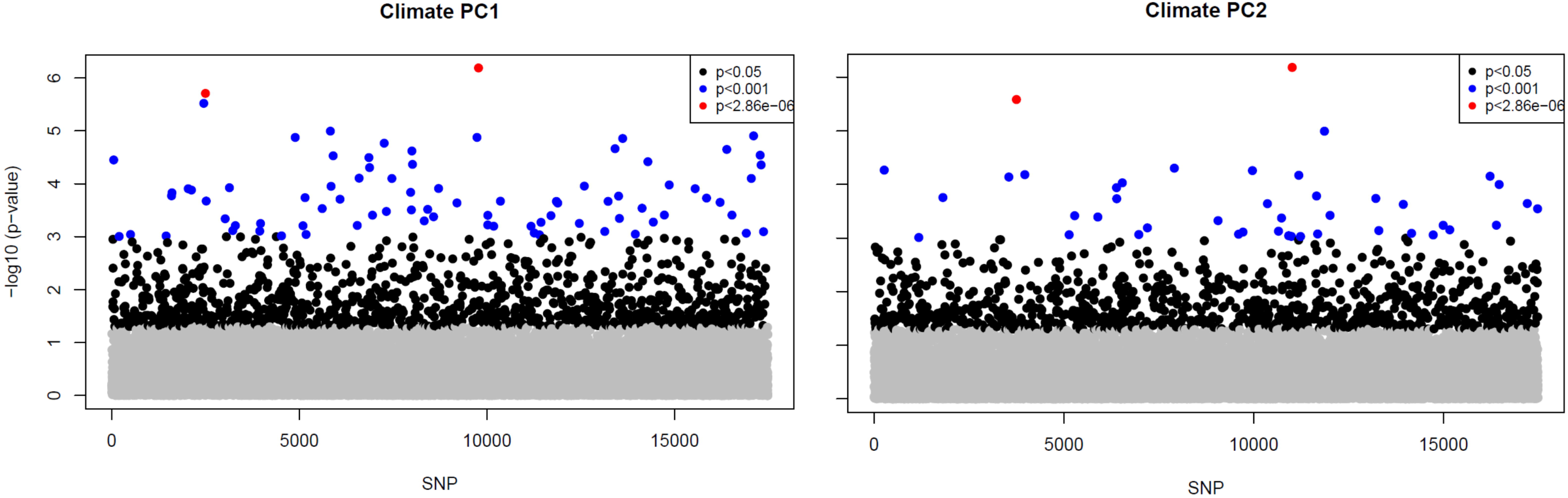

### 3.3 Outlier SNPs and environmental associations

The total number of outlier SNPs and those which showed associations with climate varied significantly among detection methods. Arlequin and LFMM2 found a total of 1148 and 1082 SNPs which showed signals of selection and association to climate PC1, respectively. However, when corrected for multiple comparisons, only 29 and 2 SNPS, respectively, passed the adjusted p-value threshold (-log_10_ corrected=5.54). BayEnv2 revealed a total of 10 SNPs associated with climate PC1 and BaySCEnv exhibited 4 SNPs (Fig.3, Fig.4a & Tab.2). 194 SNPs were commonly found by Arlequin and LFMM2, 2 SNPs by LFMM2 and BayEnv2, each 1 between Arlequin and BayEnv2 and LFMM2 and BayScEnv and all four SNPs detected with BayScEnv were also listed as outliers in Arlequin. As the most promising candidates SNPs #15099 and #78509 appeared independently as outliers associated with temperature in three of the four employed programs (Fig. 5a).

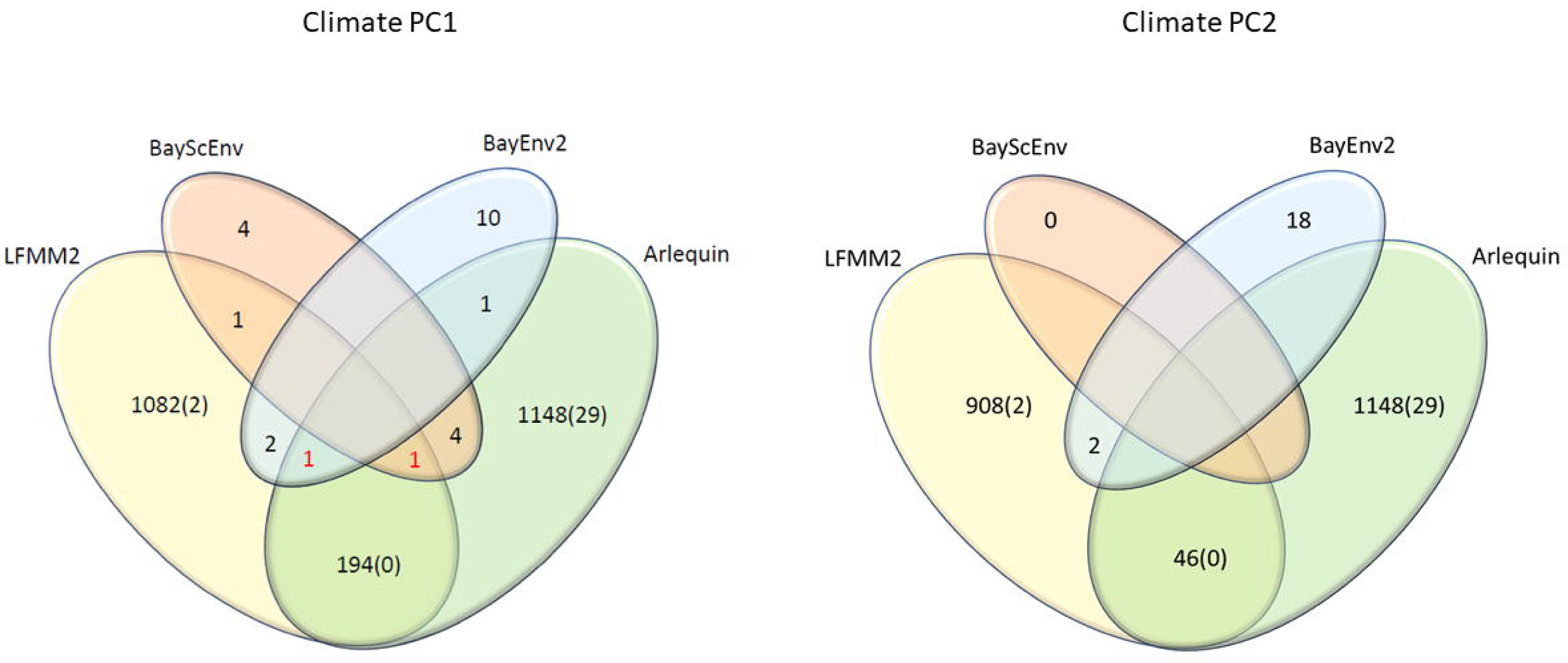

For climate PC2 LFMM2 detected 908 SNPs (2 SNPs still significant after Bonferroni correction), BayEnv2 18 SNPS while no significant SNPs were detected by BayScEnv. 46 SNPs which appeared as outliers in Arlequin were also shortlisted in LFMM2 and SNPs #51115 and #69292 were commonly deteced by LFMM2 and BayEnv2. However, for climate PC2 no SNP appeared to be shortlisted more than twice.

### 3.4 Interrogation of putative adaptive SNPs

A total of 18 consensus SNPs (11 for Climate PC1 and 7 for climate PC2) were selected for functional interrogation and spatial frequency analysis. All 18 SNPs were at least found independently by two or three of the used algorithms when the thresholds described in Table 2 were applied. However, since a large number of common SNPs were found between LFMM2 and Arlequin (194 and 46, respectively) we selected only the top five candidates ranked by the −log_10_ LFMM p-value for both climate PCs, respectively.

**Tab.1:** Overview of provenance origin and climate

| Provenance ID | Name | Region | Latitude | Longitude | MAT | MAP | N |
| --- | --- | --- | --- | --- | --- | --- | --- |
| DA | D'Arcy | British Columbia transition | 50.5567 | -122.5 | 8.3 | 516 | 13 |
| NS | Nelson | British Columbia inland | 49.5 | -117.2667 | 6.5 | 941 | 13 |
| AB | Alberni | British Columbia coastal | 49.325 | -124.85 | 9.2 | 2161 | 10 |
| NP | Newport | Washington inland | 48.2 | -117.05 | 7.3 | 736 | 11 |
| SK | Spokane | Washington inland | 47.7833 | -117.2 | 8.4 | 518 | 10 |
| ML | Matlock | Washington coastal | 47.25 | -123.4167 | 10.1 | 2353 | 11 |
| CE | Cle Elum | Washington coastal | 47.2167 | -121.1167 | 7 | 992 | 9 |
| PG | Pine Grove | Oregon coastal | 45.1 | -121.3833 | 8.5 | 453 | 10 |
| FJ | Fort St. James | British Columbia inland | 54.4833 | -124.25 | 2 | 590 | 10 |
| CM | Clemina | British Columbia inland | 52.5833 | -119.3167 | 3.9 | 854 | 9 |
| RD | Randle | Washington coastal | 46.4847 | -121.9491 | 9.3 | 1604 | 13 |
| DR | Darrington | Washington coastal | 48.1601 | -121.4968 | 6 | 3623 | 15 |
| AL | Adams Lake | British Columbia inland | 50.9651 | -119.7056 | 6.5 | 528 | 11 |
| AQ | Abiqua Basin | Oregon coastal | 44.8802 | -122.506 | 8.3 | 2191 | 11 |
| CC | Cascadia | Oregon coastal | 44.439 | -122.41 | 10.2 | 1947 | 10 |
| SP | Snoqualmie Pass | Washington coastal | 47.4129 | -121.4411 | 6.1 | 2551 | 12 |
|  |  |  |  |  |  | total | **178** |

**Tab. 2:**
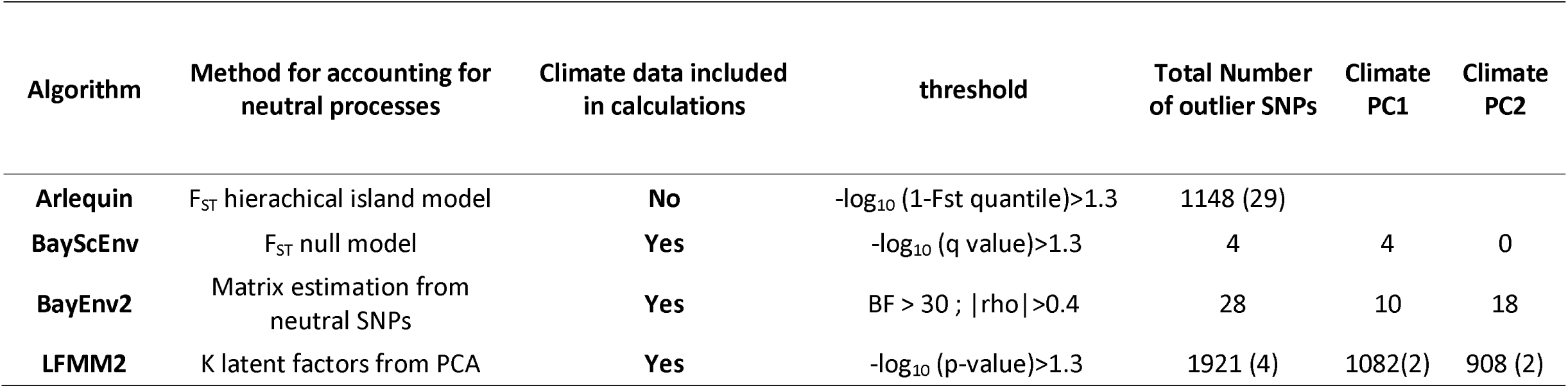
Utilized algorithms and chosen thresholds for SNP identification

For climate PC1 the analysed flanking sequences matched to *Picea abies* homologs with expect values ranging between 0 and 5.13e-06 (note that the lower the E-value the better the match). 8 out of the eleven found homologs had entries for annotated genes and for 5 of these genes putative functions were found in the congenie database (Table 3). Among those were transcription factors and proteins involved in DNA and RNA binding (e.g. TF VOZ1, Heterogeneous nuclear ribonucleo R-like isoform X), but also universally conserved proteins with still unknown function such as HemK methyltransferase family member 2 (HemK protein). Most interestingly, for SNP #78509 which was independently detected three times as outlier and significantly associated with temperature, a circadian clock protein involved in photoperiod sensing confirmed in *Arabidopsis thaliana* and *Populus trichocarpa* was found to be coded by a gene located on the *P. abies* homolog contig MA_6946339 (Tab. 3).

**Tab. 3:** Genomic positions, homolog contigs at the *P. abies* 1.0 (complete) genome assembly, and functionality of adaptive candidate SNPs. SNPs in bold were detected by three of the four algorithms.

| SNP-ID | F<sub>ST</sub> | Environmental factor (Climate PC) | Contig | Position | Ref | Alt | *Picea abies* homolog contig | E-value | Identity [%] | Function |
| --- | --- | --- | --- | --- | --- | --- | --- | --- | --- | --- |
| 15099 | **0.297343** | Temperature | LPNX010200061.1 | **392557** | T | C | MA_10430533 | **8.12E-116** | **88.15** | no entry |
| 45651 | 0.176026 | Temperature | LPNX010610246.1 | 234874 | A | T | MA_31242 | 8.68E-11 | 87.84 | Nuclear factor related to kappa-B-binding |
| **78509** | **0.412301** | Temperature | **LPNX011078866.1** | **83694** | **T** | **C** | **MA_6946339** | **3.39E-155** | **89.28** | **WD repeat-containing LWD1** |
| 20020 | 0.311698 | Temperature | LPNX010261061.1 | 122684 | G | T | MA_10426649 | 5.13E-06 | 90.00 | no entry |
| 13432 | 0.318517 | Temperature | LPNX010175959.1 | 196183 | G | T | MA_16875 | 2.23E-122 | 86.44 | unknown |
| 71701 | 0.388778 | Temperature | LPNX010985912.1 | 451465 | C | T | MA_10427644 | 8.64E-11 | 92.59 | Transcription factor VOZ1; Glycosyl transferase family 2;Heterogeneous nuclear ribonucleo R-like isoform X1 |
| 12985 | 0.364605 | Temperature | LPNX010167952.1 | 204395 | T | C | MA_14317 | 1.83E-33 | 84.10 | 60S ribosomal |
| 30680 | 0.285955 | Temperature | LPNX010408536.1 | 201415 | C | A | MA_248133 | 7.32E-16 | 90.03 | unknown |
| 70743 | 0.359548 | Temperature | LPNX010971660.1 | 402162 | A | G | MA_10428327 | 0.00E+00 | 96.10 | HemK methyltransferase family member 2 |
| 69642 | 0.303341 | Temperature | LPNX010958373.1 | 129773 | T | G | MA_10434143 | 3.48E-84 | 90.42 | unnamed product |
| 89806 | 0.30836 | Temperature | LPNX011224705.1 | 67931 | A | T | MA_206051 | 2.10E-08 | 91.23 | no entry |
| 51115 | 0.205466 | Precipitation | LPNX010683611.1 | 224568 | C | A | MA_10428611 | 7.48E-36 | 94.23 | Bifunctional UDP-glucose 4-epimerase and UDP-xylose 4-epimerase 1 |
| 69292 | 0.281179 | Precipitation | LPNX010953873.1 | 35635 | G | C | MA_2566561 | 1.70E-27 | 87.69 | no entry |
| 58318 | 0.339572 | Precipitation | LPNX010792684.1 | 264816 | A | G | MA_57244 | 2.23E-159 | 88.48 | Transcription factor FAMA isoform X2 |
| 31046 | 0.39123 | Precipitation | LPNX010411427.1 | 510083 | T | C | MA_10430971 | 1.31E-23 | 88.72 | unknown |
| 57319 | 0.265295 | Precipitation | LPNX010776984.1 | 676889 | G | A | MA_113422 | 1.78E-141 | 88.84 | no entry |
| 17435 | 0.261599 | Precipitation | LPNX010232549.1 | 270839 | T | A | MA_70676 | 6.53E-61 | 85.91 | Chloroplastic-like |
| 60539 | 0.379081 | Precipitation | LPNX010822297.1 | 383798 | G | C | MA_10430923 | 3.17E-04 | 92.31 | Exosome complex component RRP41-like |

For climate PC2 five of the seven sequences showed homolog contigs with E-values between 2.23e-159 to 3.17e-04 and for four homologs genes with putative protein functions were listed. As such, we found proteins involved in carbohydrate metabolism (Bifunctional UDP-glucose 4-epimerase and UDP-xylose 4-epimerase 1) as well as transcription factors known to be involved in stomata formation in *Nicotiana tomentosiformis* (Transcription factor FAMA isoform X2).

Finally, our two candidate SNPs #15099 and #78509, which were three times independently identified as truly adaptive outliers showed significant differentiation in space, since the alternative allele increased in frequency for both SNPs towards the inland and also towards colder provenance locations (Pearson correlation coefficients between allele frequencies and mean warmest month temperature were −0.31 and −0.32, respectively; Fig. 6).

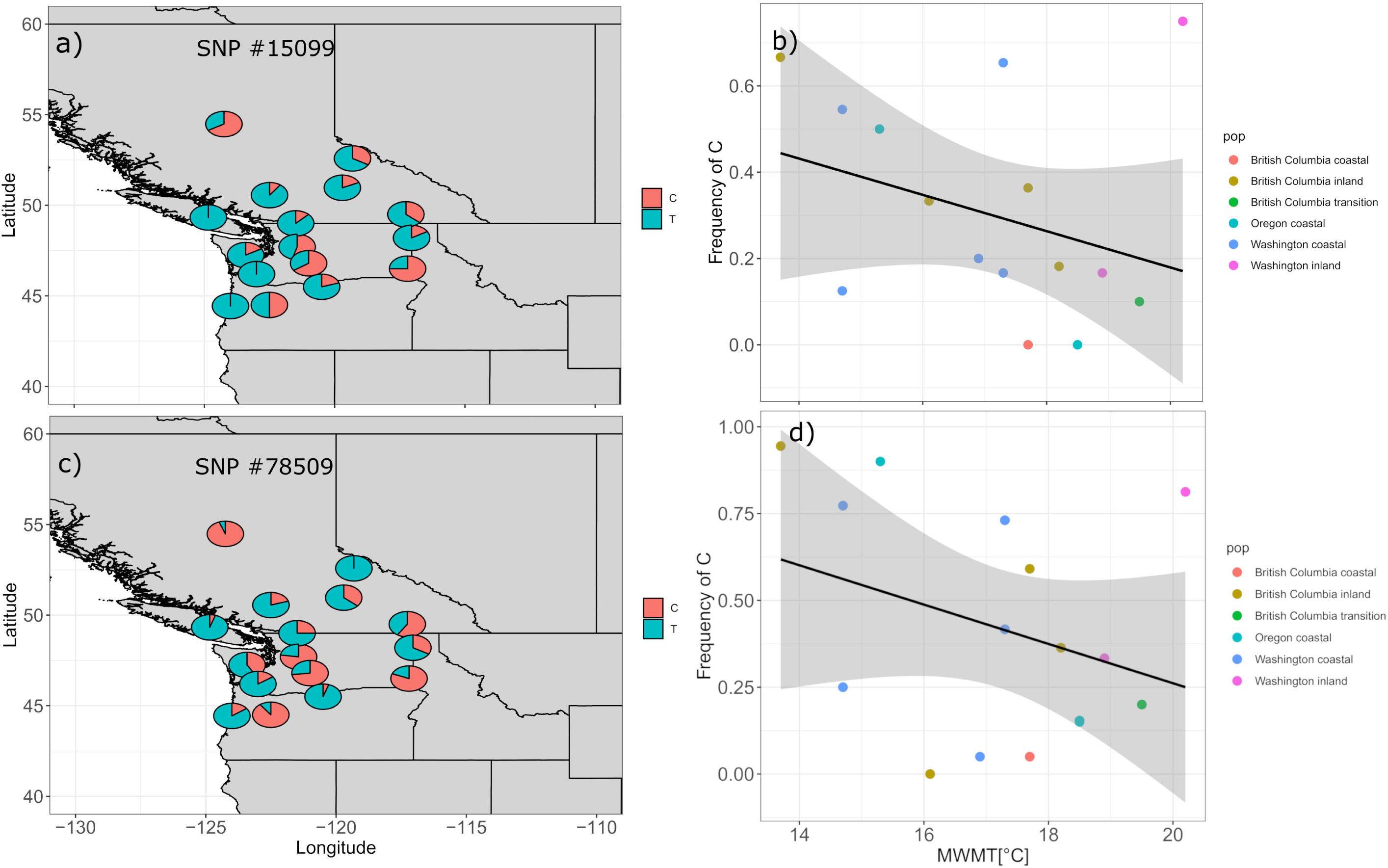

When comparing the first two eigenvectors based on the 18 consensus SNPs to eigenvectors obtained from an equal number of randomly chosen neutral SNPs, provenances could be clearly separated into inland and coastal provenances whereas no clear differentiation was observed for the neutral subset (Fig. 7).

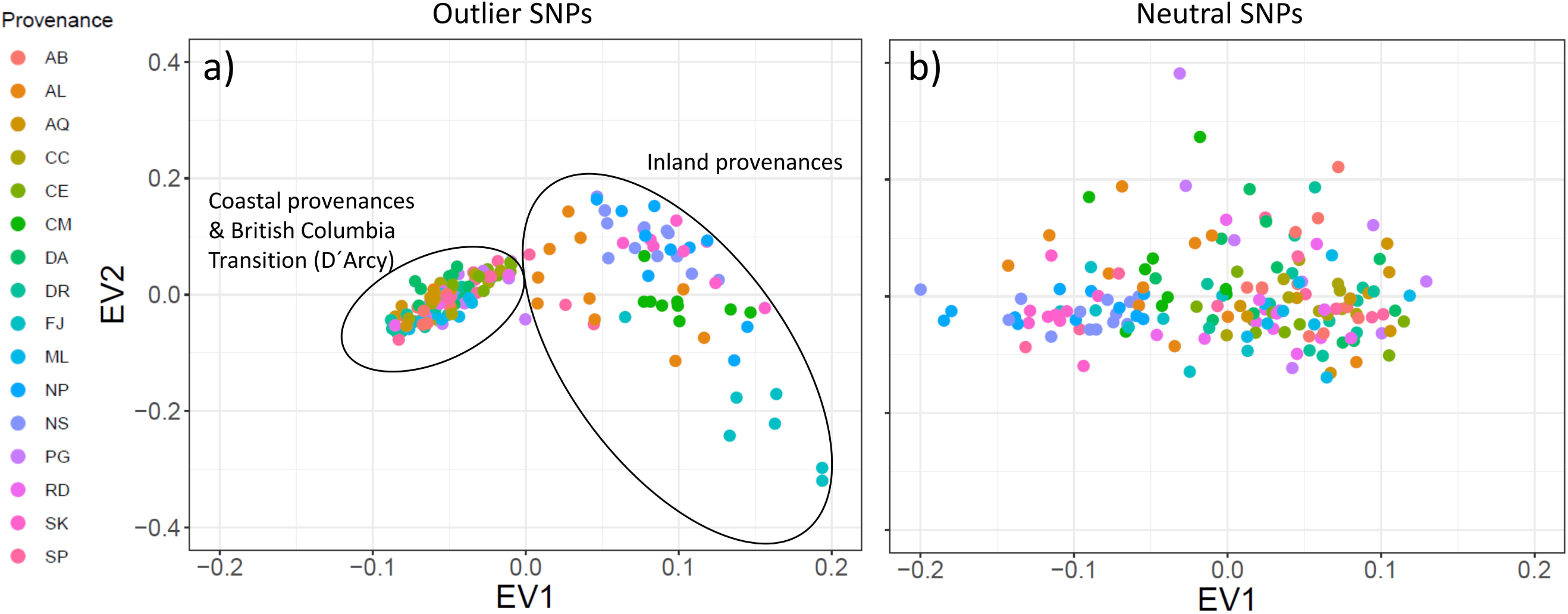

## 4. Discussion

In this study we combined observations from a common garden experiment with whole-genome information in order to unravel polymorphisms which could have caused the strong genecological cline provenances exhibited after 40 years of growing. While such old common garden experiments are rare for most tree species, their retrospective analysis by using dendroclimatology can shed light in evolutionary history and adaptation patterns, because a high number of repeated observations is naturally archived in tree-rings. Although this study is not the first attempt that linked molecular genomics with tree-ring data (e.g. Trujillo-Moya et al. 2018; Housset et al. 2018; Heer et al. 2018), it is one of the very first environmental association analyses for the economically important Douglas-fir, which is recently attracting attention outside its natural range due to its ability to cope with warming and drought (Eilmann & Rigling 2012). Hence, we will discuss our findings also in the light of adaptive forest managment.

### 4.1 Growth response of trees after four decades in the common garden

The sixteen analyzed provenances showed a strong phenotypic cline with temperature and precipitation at seed origin. Provenances with earlywood growth strongly responding to July temperature at trial site originated mainly from the warmer and wetter coastal sites in British Columbia, Washington and Oregon, whereas less responsive provenances came from colder and drier inland sites. Many other studies have revelaed such a differentiation pattern between the coastal variety *menziesii* and inland variety *glauca* either in terms of productivity (Eilmann et al. 2013), physiology (Anekonda et al. 2004), and even stress response (Kleiber et al. 2017). Our findings could indeed add evidence that coastal varieties are generally more productive, since earlywood response to summer temperature is a strong indicator for overall productivity. Earlywood cells in conifers mainly ensure that water demand in the crown will be sufficiently covered during phases of high evapotranspiration and trees with larger earlywood fractions can consequently allocate more carbon (Björklund et al. 2017). The climate at our trial site already represented the dry margin of the natural distribution of Douglas-fir (George et al. 2019) and we can assume that trees have been growing under stressful conditions in most of the years. The study by George et al. (2019) in which the same trees have been analyzed also demonstrated that provenances from warmer locations were the most tolerant ones during years with extreme water deficit, whereas colder inland provenances suffered most. However, it has yet to be confirmed whether the higher drought tolerance of coastal provenances (expressed as ratio between growth during the drought year compared to a pre-drought period) really mirrors the ability of trees to withstand drought or simply exhibits conservative growth patterns which could come at the cost of lower recovery or higher mortality after drought periods (e.g. Montwe et al. 2015).

### 4.2 Outlier SNPs and the role of selection across a steep environmental cline

One of the main goals of this study was to identify polymorphisms which show true signals of adaptation to climate and which are not confounded by neutral processes such as gene flow, drift, and migration. The analysed provenances showed strong genetic substructuring which partly concided with phenotypic differentation. This makes disentangling adaptive from non-adaptive signals particularly challenging (Ahrens et al. 2018; Nadeau et al. 2016; Rellstab et al. 2015). However, we employed four conceptully different algorithms which have implemented various methodologies for accounting for neutral processes and each of them identified a certain number of outliers. The genome scan method in Arlequin identified the highest number of SNPs, which is not surprising, since Arlequin does neither specifically associate polymorphisms to climatic factors nor does it take environmental differentiation into account. Hence, the identified SNPs are probably loaded with adaptive signals from other environmental drivers than temperature and precipitation. Surprisingly, after correcting for multiple testing only a very small number of SNPs was still standing out as truly adaptive and represented only between 0.01% (LFMM) to 0.17% (Arlequin) of the analyzed SNPs. Although direct comparison with other studies is complicated due to varying experimental settings and sample size (see for example Ahrens et al. 2018), adaptive SNPs in other genera such as *Fagus* (Pluess et al. 2015), *Populus* (Fahrenkrog et al. 2017), *Alnus* (DeKort et al. 2014), and *Pinus* (Ruiz Daniels et al. 2019) comprised between 1.6% (Fahrenkrog) to 11% (Pluess et al. 2015) of all SNPs that were analyzed. Moreover, when the data was further aggregated to consensus SNPs which were simultanously detected by more than one method, only 11 candidates out of 17489 (0.06%) were detected for temperature and 7 (0.04%) for precipitation regimes. Since the analyzed provenances exhibit a strong patterm of population substructuring into five clearly delimited clusters, we strongly presume that this neutral variation is the most important cause for this finding. In particular, provenances D’Arcy (BC) and Snowqualmie Pass (WA) represented strongly isolated subpopulations based on their neutral genetic pattern (Fig. 1c), although their phenotypic signal fit well into the observed environmental cline with both climate PCs (Fig 2).

The 18 consenus outlier SNPs that were either associated with temperature or precipitation were capable of discriminating between two clusters of provenances which could be clearly assigned to one coastal and one inland group (Fig. 7). This underpins that these SNPs have most likely featured phenotypic and genetic differentiation in Douglas-fir beyond neutral processes such as isolation-by-distance. Interestingly, provenances D’Arcy (BC) and Snowqualmie Pass (WA) appeared not any longer as single subpopulation clusters when the outlier dataset was used for clustering (compare to Fig. 1c with the neutral dataset of 1,500 SNPs). One possible explanation could be that isolation of these provenances occurred relatively late compared to climatic adaptation. This finding would be corroborated by the fact that both populations showed a rather expected phenotypic signal which was similar as for the other coastal provenances (Fig. 2). Coastal and interior Douglas-fir varieties have most likely diverged during orogeny of the Cascade range, which have led to xerification of the Great Basin during the Pliocene around 2 Ma ago (Gugger et al. 2010; Brunsfeld et al. 2001). However, population contraction and expansion around the last LGM 18ka ago could have caused local spots with limited gene exchange, in particular in areas with high mountain barriers (Gugger & Sugita, 2010).

Nevertheless and despite the overall small number of SNPs that was found, we were able to extract two hot candidate SNPs associated with temperature regime at seed origin. Minor allele frequencies of both SNPs showed significant correlation with mean warmest month temperature in space, which is highly correlated with the initial response climate variable from the common garden (correlation of July temperature and MWMT=0.99). In both cases, the frequency of the minor C allele increased towards inland areas with lower temperature. In light of the strict filtering applied in this study and given the number of employed algorithms, the evidence strongly suggests that these SNPs could have featured adaptation to temperature regimes in Douglas-fir across the steep gradient that was analyzed here. Although the correlation between allele frequencies and temperature regime was moderate and characterized by a few unexpected outliers, it is likely that the rather small sample size of analyzed trees within provenances can be responsible for that. We hence see this result as a starting point for further investigations including also landscape genomics in order to corroborate these findings with more data.

### 4.3 Putative biological functions of candidate SNPs

Many of the consensus SNPs had homolog contigs at the annotated *Picea abies* reference genome from Nystedt et al. (2013) and we found some contigs containing genes with an annotated known biological function (Tab. 3). While not all of them refer directly to functions related to temperature response or water supply, their further interrogation could be though valuable, since some of the transcription factors and proteins are, for instance, involved in carbohydrate metabolism. For example, Bifunctional UDP-glucose 4-epimerase and UDP-xylose 4-epimerase 1 are important catalysator proteins which are involved in the process of cell wall biosynthesis and cell wall organization in stem tissue of higher plants (see for more information the Uniprot entry Q42605 under https://www.uniprot.org) which in turn is an important trait in softwood species confering drought resistance (Isaac-Renton et al. 2018).

Most interestingly, the triple-found SNP #78509 had a highly similar homolog contig which contained one annotated gene coding a WD repeat-containing LWD1 protein. This protein belongs to the circadian clock protein family and is regulating circadian period length and photoperiodic flowering in *Arabidopsis spec*. (Wang et al. 2011; Wu et al. 2008). Moreover, a closely related protein of this family, named GIGANTEA-5, is involved in adaptation to temperature regimes (Cao et al. 2005) and was also found to be under strong selection in *Populus balsamifera* (Keller et al. 2012; Fitzpatrick & Keller 2015). In the study by Fitzpatrick & Keller (2013), turnover in frequency of SNPs located in this gene were best explained by differences in temperature regimes among provenance origins, which strongly corroborates the findings of this study. We see this finding hence as a putative needle in the haystack, which could be the starting point for further investigations at landscape level.

### 4.4 Douglas-fir genomic resources for adaptive forest management

Douglas-fir is currently discussed as a promising substitute species outside its natural range given its putatively higher drought tolerance and yield (Chakraborty et al. 2016; Eilmann & Rigling, 2012). Our results could be pertinent for future studies and applications which are aiming at identifying adapted seed sources for Douglas-fir. For instance, SNP information can be used in order to genotype large arrays of trees from progeny trials or provenance tests in order to decide whether the selected material is generally site-adapted or likely to be maladapted to temperature and drought regime. Additionally, our dataset can be used as reference dataset for genomic assignment of individuals or stands given that many Douglas-fir seed stands in Europe are still without confirmed geographic origin. Therefore, we provide detailed SNP and phenotypic information such as genomic positions, flanking sequences, and rank scores for more than 17,000 SNPs obtained from the various programs in the Supplementary Material S2.

## 5. Conclusions

Based on a steep phenotypic cline observed in a common garden experiment we were able to disentangle adaptive signals in Douglas-fir from those that were simply caused by neutral demographic processes. We showed that combining dendroclimatological data with whole-exome information can lead to valuable insights into the adaptation history of a widespread conifer. Although a very small fraction of the analyzed polymorphisms stood out as adaptive candidates, their functional interrogation strongly suggests that SNPs could have indeed featured climatic adaptation in Douglas-fir. Thereby, we shed new light into the adaptive history of another conifer with high economic and ecological importance.

## 6. Acknowledgements

This study is part of the Translational Research Program 122 (‘Softwood for the Future’) and was funded by the Austrian Science Fund (FWF). Dr. Marcela van Loo was funded by the Austrian Science Fund (FWF) project „Verhalten der ‘Europäischen’ Douglasie” (P26504). We also would like to thank the Austrian Central Institute for Meteorology and Geodynamics (ZAMG) for providing weather data as well as Michael Eberhardt for help during the field work. We are also grateful to Thorsten Günther (Uppsala University) for advice with Bayenv2.

